# Asian pika populations track local glaciation events through the Pleistocene

**DOI:** 10.1101/2025.06.25.661461

**Authors:** Harsha Kumar, Nagarjun Vijay, R Nandini

## Abstract

**Background:** The Pleistocene glaciation cycles (2.6 mya - 11 kya) were major climatic events that shaped diversity and community assemblages on Earth. Cold-adapted, high-elevation specialists are expected to have responded negatively to warm interglacial periods, with populations contracting and being pushed up the elevational gradient, leading to isolated populations on mountain tops. During the cool glacial maximums, they are expected to have been distributed at both low and high elevations, with populations expanding and facilitating gene flow across mountains. Pikas (*Ochotonidae*) are poorly studied high-elevation specialist Lagomorphs being extirpated at alarming rates due to climate change. Insights into their historical demography and current effective population size are crucial in determining their current population status to inform conservation planning.

**Results:** We use de novo assembly to construct partial genomes of three Asian pika species (*Ochotona ladacensis, O. nubrica,* and *O. macrotis*) that differ in their life history, lifestyle, and social behaviors. We use a combination of species distribution modeling (SDM) projected back in time, in addition to inferring historical demographic patterns using a pairwise sequentially Markovian coalescent (PSMC) approach. Our SDMs predicted the largest bio-climatic niches for all three species during the last glacial maximum (LGM ∼20 kya), and the smallest niches at the last interglacial (LIG∼120 kya), agreeing well with the elevational shift hypothesis. Our PSMC models for the populations in the Changthang Biotic province (CBP), revealed largely synchronous oscillations of populations sampled in geological timescales but displayed contrasting temporal patterns of population spikes when compared with SDMs. The largest effective population sizes of all species examined on the CBP were inferred to pre-date both the LGM and LIG and were placed around the MIS-6 (∼ 191 kya). After the MIS-6 local glaciation event, populations were inferred to have been through a deep bottleneck, correlating well with local glaciation patterns on this plateau.

**Conclusion:** Our study illustrates the power of using complementary approaches to infer historical demography in mountainous landscapes with complex glaciation histories. While SDMs appropriately describe species responses to historical climate across the geographic range of species, PSMCs are more informative about local population dynamics when subject to large degrees of population differentiation. All three species of pikas responded similarly to historical glaciation events on the CBP that differ from the rest of the world and other parts of the Himalayas and likely have very low effective population sizes at current timescales. This is one of the first few studies to examine the population demography of small mammals on the Indo-Tibetan Plateau and has the potential to inform species and population-specific conservation planning in the Himalayas. It is very likely that small mammals such as pikas, marmots and voles on the CBP have low effective population sizes due to historical inbreeding during the LIG and the lack of a glaciation rescue during the LGM. This highlights the need for conservation measures for small mammals on the landscape.

## Background

Climate change-associated warming and erratic weather patterns are expected to cause rapid declines in biodiversity worldwide [1, 2]. Alpine specialists on mountain tops are at differentially higher risk of extinction, given their limited potential to shift ranges [3–5]. Since most species living in remote landscapes have not been assessed for extinction risk, identifying traits that make animals more susceptible to climate change is useful for species management [6–9]. Species at risk today have survived large-scale climatic events during the Pleistocene glacial cycles, and gaining insights into their historical responses might help us predict how they respond to future climate change [10]. Pikas of the family *Ochotonidae* are a unique lineage of small mammals adapted to life on high-elevation cold mountains [11, 12], and have been recognized as ecologically important sentinels that are under threat due to climate change [13, 14].

Species distribution modeling (SDM) approaches offer insights into how species might have responded to historical climate change [15, 16]. By understanding ecological factors that govern species distributions today, we can project preferences of species into the past or the future, assuming niche conservatism over evolutionary time scales [17–21]. SDMs in their simplest form don’t account for critical biotic interactions that govern community structuring [22–24] and work best at shallow timescales, given that species can adapt to a changing climate over large evolutionary timescales [25] .

Recent advancements in population genetics and coalescent theory have allowed researchers to capture genomic signatures of historical demographic fluctuations [26–30]. The pairwise sequentially Markovian coalescent (PSMC) analysis can work on a single genome by using nucleotide variation between two haploids to reconstruct demography [27, 30]. It formalizes the relationship between the rate of coalescence of alleles at different loci and effective population size under a model of neutral evolution [27, 30]. Although PSMCs are sensitive to selection effects, population structure, and the inclusion of mitochondrial/sex chromosomes; they can be reliably used to understand deep timescale (20kya-1000kya) demography when interpreted with caution [27, 30–34]. Given that the two methods work optimally at different temporal and spatial scales (historic vs current and populations vs species), complementing PSMCs with SDMs allows researchers to understand how species have responded to climate change historically and how they might meet climate change in the Anthropocene [35–38].

The Indo-Tibetan plateau is a natural laboratory for the study of adaptations and species responses to complex glaciation patterns [39]. Its high elevation, extreme climate and dynamic geological history have challenged species to adapt or migrate to more favourable climates. While many researchers have studied demography on the plateau using species distribution models across geological timescales, it is focused heavily on plants [21, 40–44]. Inferences of historical demography of species using population genomics from the region however, has been greatly limited to large mammals [45–49]. While there are a handful of studies on other taxa such on small mammals such as pikas[50], birds [36, 51], plants [35] and, butterflies [52]; there is scope for a lot more work on the landscape given that glaciation histories are vastly different across this region [43, 53–57].

Our understanding of pika biogeography has large implications for the conservation of these specialists [12]. Pika populations are expected to track cool habitats, move up the elevational gradients at warm time points in the Earth’s history, and expand to lower elevations at cool time points (Elevational shift hypothesis) [58] (Figure 1). This hypothesis is supported by the: a) Eocene pika fossil record: Specimens found exclusively at high elevations (Qinghai-Tibetan Plateau, the Alps, and the Rocky Mountains) corresponding to a warming event [12, 59]; b) Miocene - Pliocene fossil record: Specimens found at both low and high elevations corresponding to a cooling event [59]; c) Pleistocene fossil records: Specimens of *O. princeps* found at low elevations corresponding to glacial maximums [58]. In this study, we use a two-pronged approach (SDMs and PSMCs) to examine the historical demographic responses of three sympatric species of cold-adapted mountain specialist pikas from the Changthang biotic province of the Tibetan plateau. *Ochotona ladacensis* is a social, burrowing, high elevation specialist (>4200 m) occupying alpine meadows*, O. nubrica* is a social, burrowing species that utilizes a large elevational gradient (3300-6000 m) and is an alpine scrub specialist [60]*. Ochotona macrotis* is a rock-dwelling, asocial species that also occupies a large elevational gradient [60]. Given that pikas are cold adapted specialists, we expect that: a) population sizes and fundamental niches of all three species would be smallest at the last interglacial (LIG; 120kya) and largest at the last glacial maximum (LGM; 20kya) (Hypothesis 1); b) burrowing species would be more resilient to historical climate change when compared to rock-dwelling species since burrows provide better insulation against extreme weather when compared to rock refuges [98] (Hypothesis 2); c) *O. ladacensis* that exclusively occupies high elevations (>4500m) at current timescales, is likely to have small effective population sizes (Hypothesis 3). This is due to warming events causing severe bottlenecks, driving populations up the elevational gradient much more than they would for species that have wider elevational distribution. Understanding how small mammals from the Tibetan plateau have responded to historical climate change will likely be useful for managing alpine specialists.

**Figure 1:**
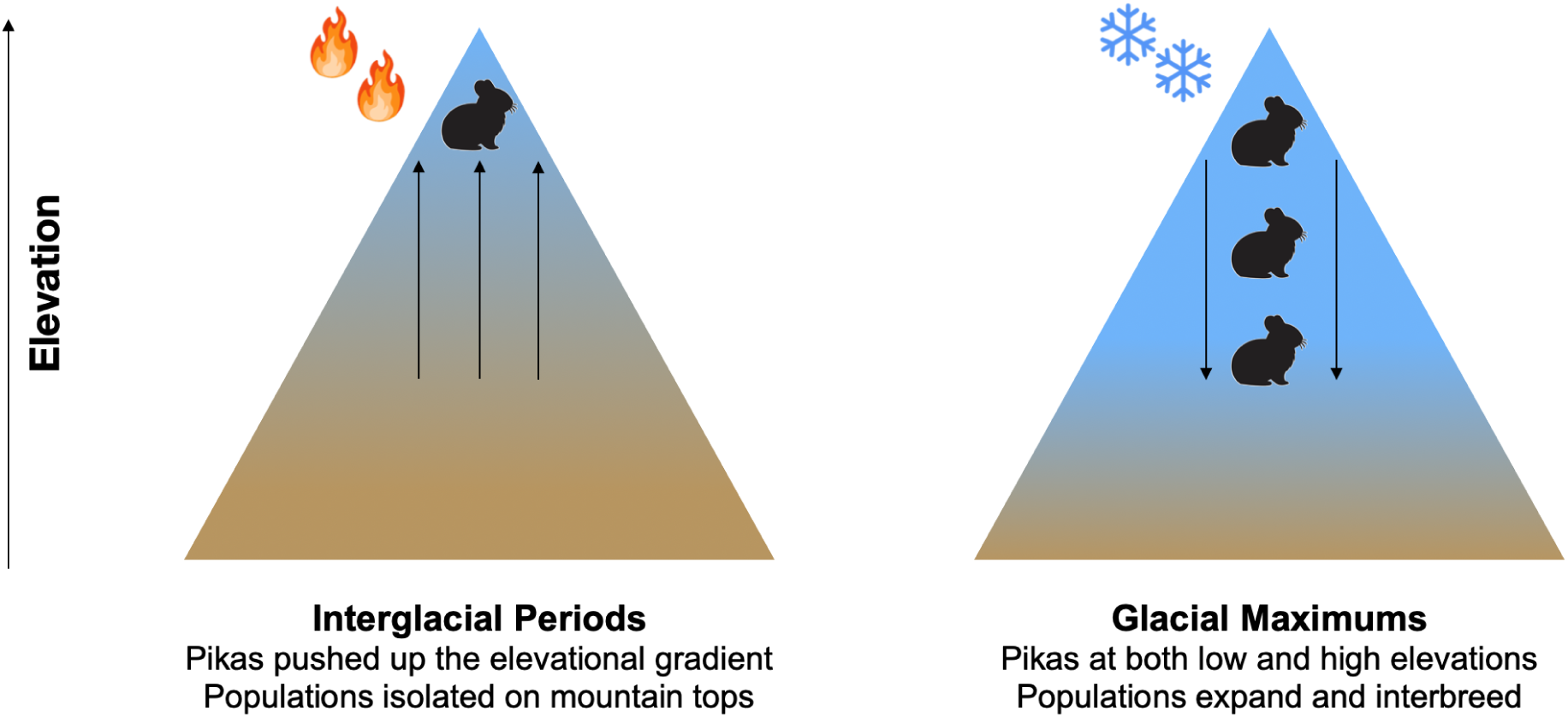
The elevational shift hypothesis describing how pika populations track cool climates during the Pleistocene glacial cycles (adapted from [58]).

## Results

### Draft genome assemblies for pikas

Our genome completeness were assessed by both metrics of Megahit N5o and BUSCO. N50 measures were largest for *O. ladacensis* (25741 bp), followed by *O. nubrica* (15016 bp) and finally *O. macrotis* (N50 - 8050 bp). BUSCO scores had a large variance with 53% of complete BUSCO groups being sequenced for *O. nubrica*, 41% for *O. ladacensis* (N50 - 25741 bp) and 19% for *O. macrotis;* with 13%-18% being sequenced partially across species (Figure 2A). Our Orthovenn3 analyses indicated a large proportion of shared genes clusters across species examined (830 shared between species sequenced in this study; 445 shared between species sequenced in this study and *O. princeps*). We also found equally large proportions of gene clusters being uniquely associated with the two partial genomes of *O. nubrica* (462) and *O. ladacensis* (387) (Figure 2B).

**Figure 2:**
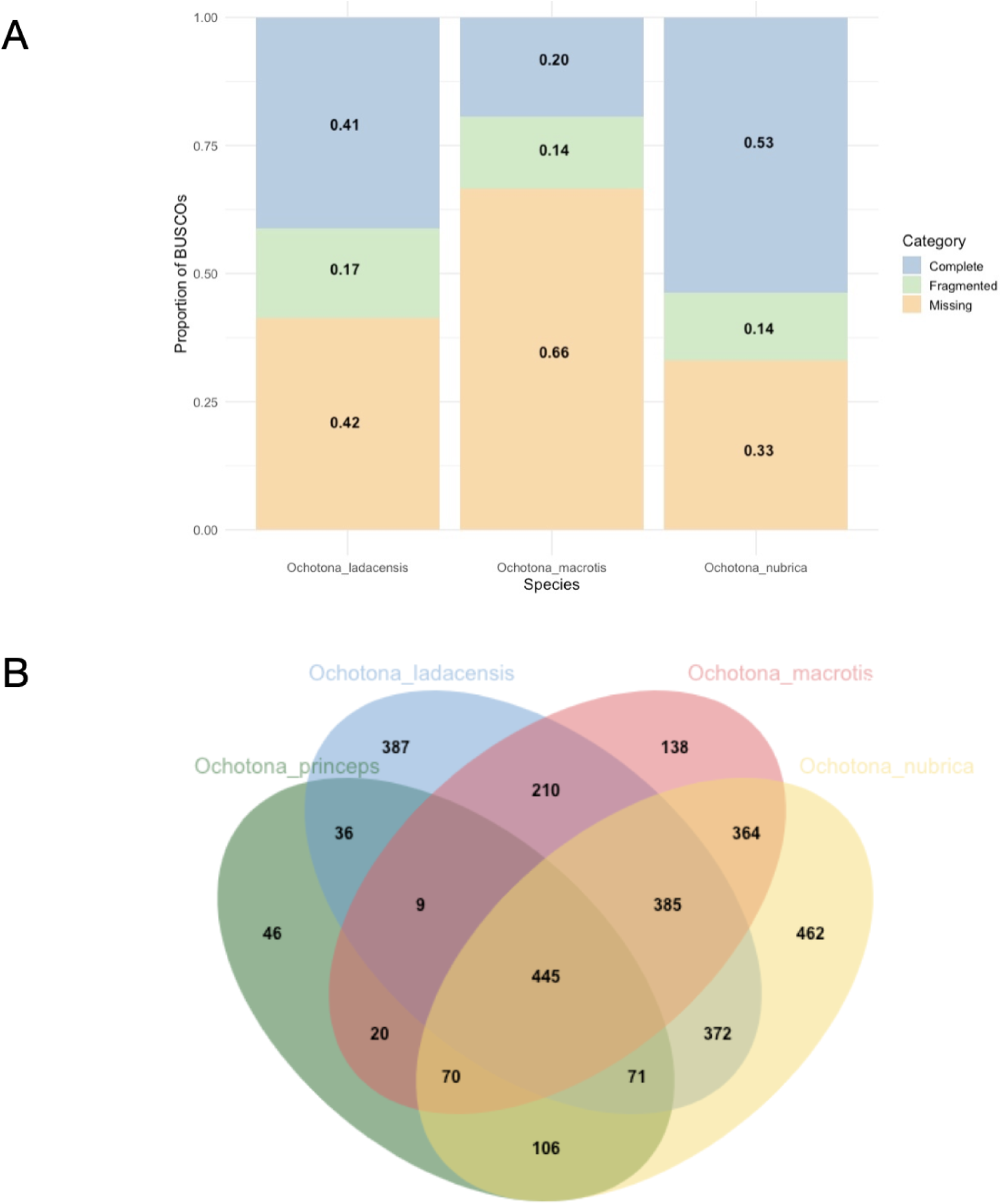
A - Proportion of BUSCO groups sequenced from the mammalian database (N=9226 BUSCO groups) across the three species. B - Orthologous gene clusters found between species using OrthoVenn3 using the protein prediction tool Augustus.

### Projecting bio-climatic niches into the past: pikas have the largest niches at LGM and the smallest niches at LIG

Ladakh pika (*Ochotona ladacensis*) - The best models constructed for the species had an AUC score of 0.967 with precipitation seasonality (Bio15), mean temperature of warmest quarter (Bio10), mean temperature of the wettest quarter (Bio8) and precipitation of the driest month (Bio 14) being the most critical variables driving the current distribution of the species (Table 1, Figure S1). Niche suitability (suitability > 0.4) for this species was inferred to be the smallest at LIG (∼4800 km^2^) and expanded significantly at LGM (a 160-fold increase). Niches for this species were inferred to have shrunk post the LGM into the mid-Holocene warming (∼ 5-fold decrease) and expanded slightly at current timescales (2 % increase since mid-Holocene) (Figure 3).

**Figure 3:**
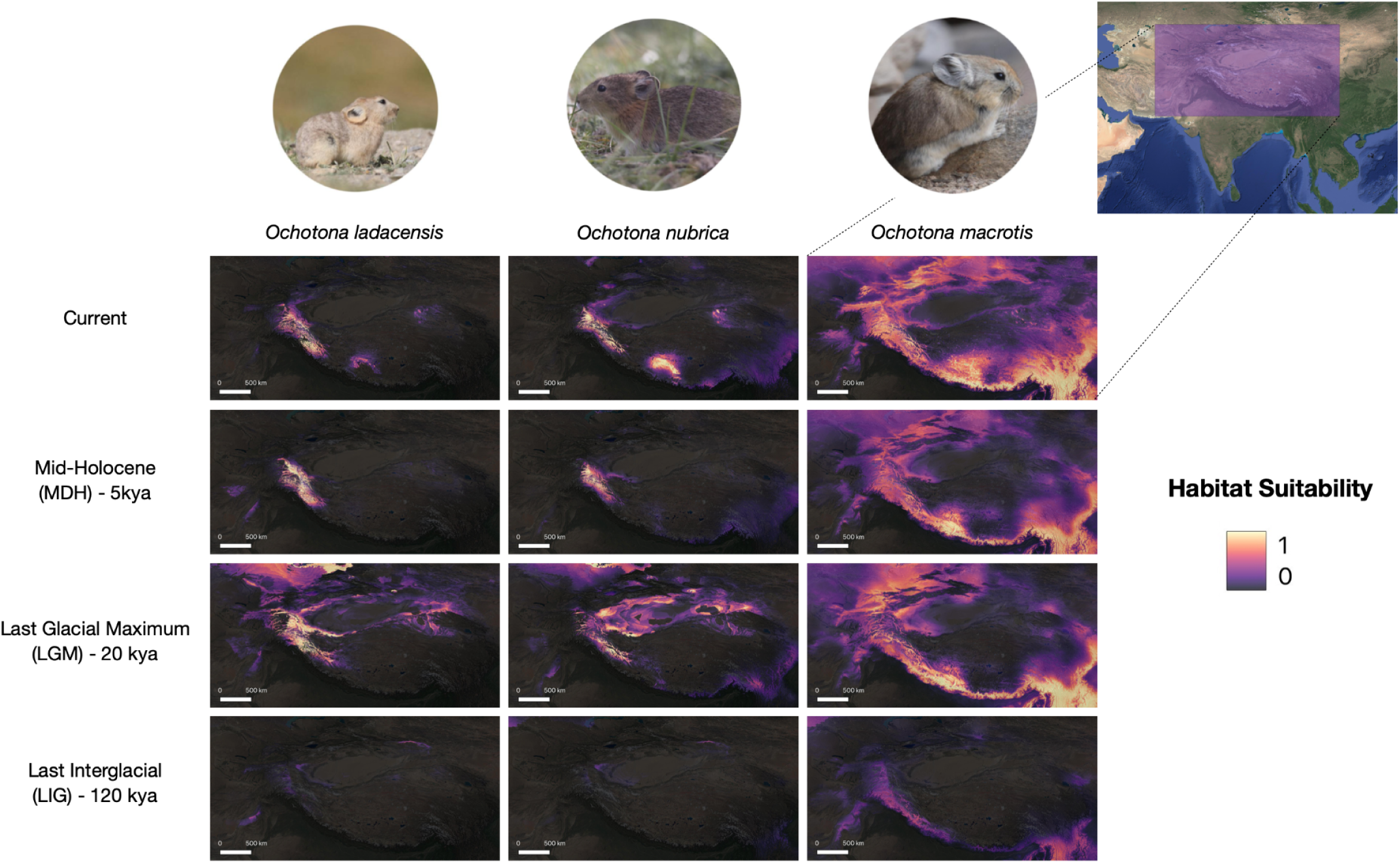
Habitat suitability maps for different species (columns) at different time slices (rows): Current, Mid-Holocene (5kya), Last Glacial Maximum (20kya), and Last Interglacial (120kya). Warmer colors indicate higher suitability of habitats for species.

**Table 1:** Top variables (Percent contribution > 5%) identified in MAXENT models of the three species of pikas.

| Species | Variable | AUC | Percent contribution | Permutation importance |
| --- | --- | --- | --- | --- |
| *Ochotona nubrica* | bio9_mean_temp_dri_qtr |  | 28.8 | 27.4 |
|  | bio15_precip_seasonality |  | 17 | 36.1 |
|  | bio7_temp_ann_rang |  | 11 | 0 |
|  | bio17_precip_dri_qtr |  | 8.6 | 7.2 |
|  | bio8_mean_temp_wet_qtr |  | 7.8 | 22.7 |
|  | bio19_precip_cold_qtr |  | 7.3 | 5.9 |
| *Ochotona ladacensis* | Variable | AUC | Percent contribution | Permutation importance |
|  | bio15_precip_seasonality |  | 29.3 | 8.4 |
|  | bio10_mean_temp_wrm_qtr |  | 26.8 | 0 |
|  | bio8_mean_temp_wet_qtr |  | 13 | 30.8 |
|  | bio14_precip_dri_mnth |  | 10.2 | 0.8 |
|  | bio3_isothermality |  | 7.8 | 13.4 |
|  | bio9_mean_temp_dri_qtr |  | 7 | 22 |
| *Ochotona macrotis* | Variable | AUC | Percent contribution | Permutation importance |
|  | bio1_ann_mean_temp |  | 26.7 | 27.7 |
|  | bio14_precip_dri_mnth |  | 18.9 | 0.6 |
|  | bio9_mean_temp_dri_qtr |  | 16.5 | 11.8 |
|  | bio15_precip_seasonality |  | 15.9 | 21.4 |
|  | bio11_mean_temp_cold_qtr |  | 6.6 | 0 |

Nubra pika (*Ochotona nubrica*) - Best models constructed for the species had an AUC score of 0.972, with the mean temperature of the driest quarter (Bio9), precipitation seasonality (Bio15) and temperature annual range (Bio7) being the most critical variables driving current distribution of the species (Table 1, Figure S1). Models projected back into the past indicated that a small area was suitable for the species at LIG (∼1600 km^2^), with a significant expansion of bio-climatic niches at LGM (a 400-fold increase in suitable habitat). Subsequently, there was a significant reduction in niches for the species during the mid-Holocene warming (82% reduction since LGM). Suitable niches have expanded post the mid-Holocene into the current timescales (a 2.5-fold increase) (Figure 3).

Large-eared pika (*Ochotona macrotis*) - Best models constructed for the species had an AUC score of 0.918 with annual mean temperature (Bio1), precipitation of driest month (Bio14), mean temperature of the driest quarter (Bio9) and precipitation seasonality (Bio15) being the most critical variables driving current distribution of the species (Table 1, Figure S1). Niche suitability for this species was inferred to be the smallest at LIG (∼ 240000 km^2^), expanded significantly at LGM (a 10-fold increase from LIG), shrunk post the LGM into the mid-Holocene (∼ 20% decrease from LGM) and expanded by a large extent at current timescales (67 % increase since mid-Holocene) (Figure 3).

Taken together, this yields support for our hypothesis that pikas have larger niches at LGM when compared to other time points (Hypothesis 1). Univariate response curves (Maxent models run only with one variable) showed that the rock-dwelling species *O. macrotis* had the largest bioclimatic envelope, which largely explains its extensive distribution today (Figure S1). In particular, *O. macrotis* presence was associated with a wide range of annual temperatures (Bio7), mean temperature in the wettest quarter (Bio8), precipitation in the driest month (Bio14), and precipitation seasonality (Bio15) when compared to *O. nubrica* and *O. ladacensis*.

### Pairwise sequential markovian coalescent (PSMC) simulations: pikas of the Indo-Tibetan plateau have the largest population sizes around the MIS-6

Estimates of effective population sizes of different species of pikas from the CBP followed consistent trends (Figure 4). We estimated the largest effective population sizes for species during a) the warm LIG (∼120kya) for *O. nubrica*, and b) the cool MIS-6 (∼191kya) for *O. macrotis and O. ladacensis* (Figure 4). Effective population sizes of all species were inferred to have continued to drop post the LIG, through the LGM (∼20kya), into the Mid-Holocene, with only *O. nubrica* showing a small recovery post the LGM (Figure 4). We find that the species that are currently restricted to high elevations; namely *O. ladacensis* has the smallest effective population sizes (Hypothesis 2). Estimates of effective population sizes were not affected by lifestyle; with the rock-dwelling species examined (*O. macrotis*) being sandwiched between the two burrowing species (*O.ladacensis*, *O.nubrica*) (Figure 4).

**Figure 4:**
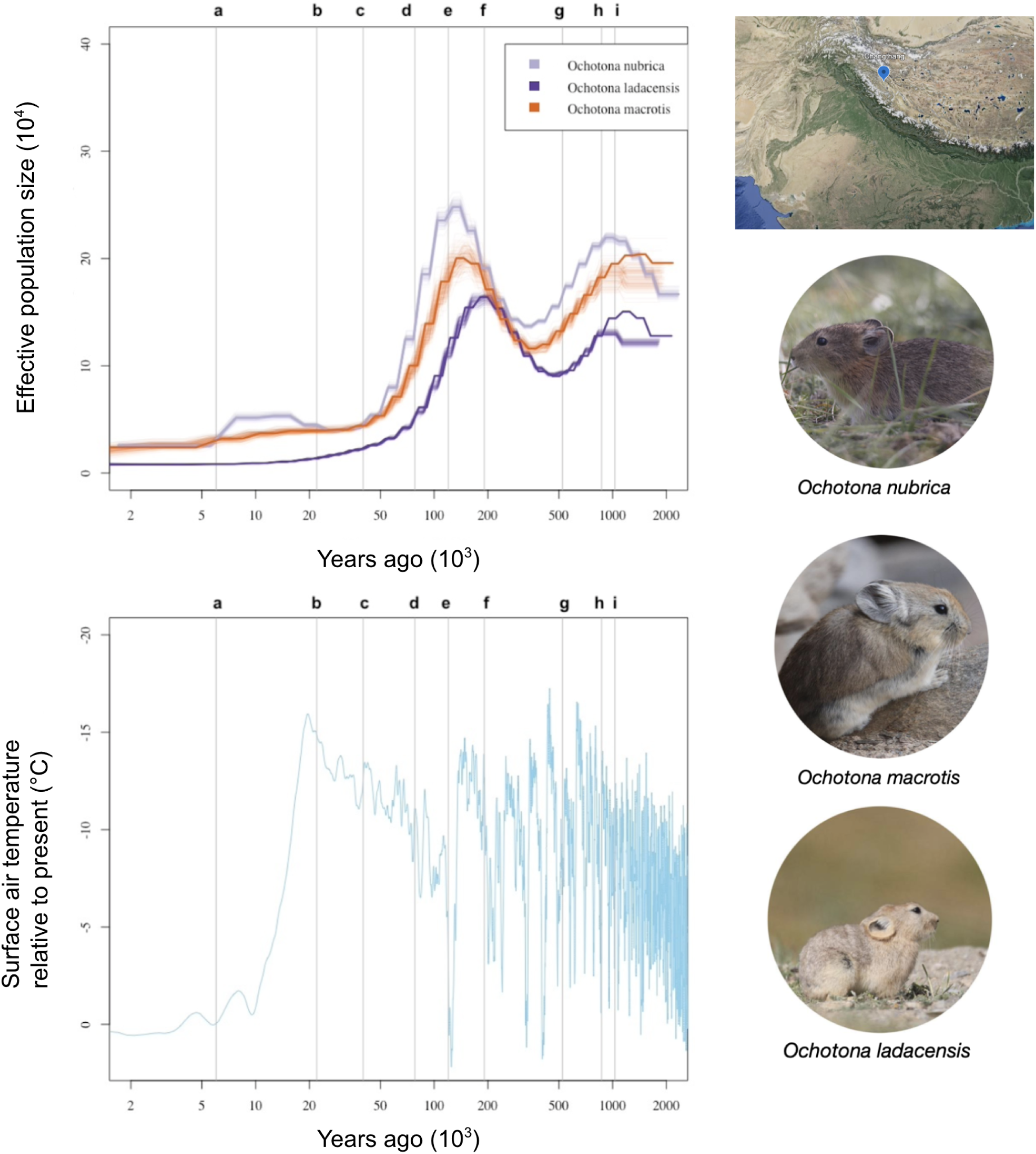
Top left - PSMC plots of three sympatric species of pika from the Changthang Biotic province showing estimates of effective population sizes between 2kya-2mya; Bottom left - Corresponding average atmospheric surface temperature relative to present constructed using GCMs (data from [97]. Legend: a - Mid-Holocene; b-Global last glacial maxima; c-d- Local last glacial maxima/Changthang Biotic province glaciation event (MIS3 - MIS4); e - Global last inter-glacial period; f - MIS-6; g - MIS-13; h - MIS-21; i - MIS-22.

## Discussion

Studying the historical demography of species in the context of climate change over evolutionary timescales provides a window into past species’ responses. In this study, we aimed to recover the demographic histories of cold-adapted high-elevational specialist pikas from the Indo-Tibetan Plateau using a combination of SDMs and PSMCs. We detected low effective population sizes for pika populations at LGM using PSMCs. This contrasts our species distribution models which suggest large geographic distributions of species at this time point. Accounting for local geology and historical climate experienced by different populations, in addition to consideration of global distribution patterns will be useful for the conservation planning of these species.

### Pikas populations move up the elevational gradient during LIG and expand to lower elevations during LGM

The SDMs predicted and supported our hypotheses that the last glacial maximum (∼20kya) was likely a time point when pika populations were large (Hypothesis 1). We recovered large niches at the LGM and small niches at LIG for ⅔ species (*O. nubrica, O. ladacensis*) (Figure 2). Habitats of moderate-high suitability (>0.4) were typically at lower elevations during the LGM when compared to current time scales following the elevational shift hypothesis [58] (Table S3). However, suitable habitats for the species *Ochotona macrotis* alone were at higher elevations during the LIG compared to the LGM (Table S3). This is potentially due to the complex glaciation history and historical climate of the Himalayas detailed below. SDMs for other cold-adapted high-elevation plants and frogs on the Tibetan plateau through the Pleistocene have revealed complex patterns involving both expansions (*Kobresia* spp, *Toxicodendron vernicifluum, Rhodiola* spp, *Allium mongolicum*) [40, 61–63] and contractions (*Kobresia* spp, *Larix sibirica*, *Nanorana pleskei, Primula* spp, *Mirabilis himalaica, Anisodus tanguticus*) [41, 44, 62, 64–67] during the LGM, linked to species ecology and thermal physiology.

### The historical climate of the Indo-Tibetan plateau and its impact on native flora

A biogeographical and geological perspective is needed to understand the population-level responses of pikas to historical climate change. The complex geology of the ongoing upliftment of the Himalayas and the expansion of the Tibetan Plateau that began in the Pliocene has caused large-scale changes in climate, landscape, and vegetation at local scales, driving the evolution of new groups of mammals [68, 69]. Our PSMCs placed the largest population sizes of species at MIS6 (∼190kya), pre-dating the LGM and LIG. This can be explained by the large disparity in the climatic records between the ‘global last glacial maxima (GLGM: 18kya to 22kya)’ experienced by the Northern Hemisphere and the ‘local last glacial maxima (LLGM: 40kya - 78kya)’ experienced by the Tibetan Plateau and the Himalayas during the ‘Batal stage’ as evidenced by geological records [43, 53–57]. The largest glacial maximum events on the Indo-Tibetan plateau, however, are known to have occurred during the late Pliocene (MIS-12 - MIS-6 strata; 0.175mya - 0.5mya) and differ from glacial dynamics from the rest of the world [70–72]. More interestingly, glaciation patterns experienced by different parts of the Indo-Tibetan plateau such as the Changthang biotic province (CBP) and the Lauhul Himalayas are also significantly different [43, 53, 54, 56, 68]. The local glacial maxima experienced by the CBP from where the genomes were sampled, has been dated to span much of the MIS-3 (57 kya) - MIS-4 (71 kya) and was characterized by extremely low precipitation (snow) and small disconnected glacial valleys (<10km from current ranges) [43, 53, 54, 56, 68] as opposed to large interconnected glacial valleys predicted through the LGM in the Northern Hemisphere [73, 74]. This pushes the timeline of the actual ‘tiny’ LLGM at our sites back to 78kya, likely lasting till 40kya [43, 53, 54, 56, 68], situating it way closer to the global last interglacial (120kya). Additionally, pollen analysis from the Tsokar Lake Basin on the CBP suggested periodic expansion of *Juniperus* communities and marshy vegetation (indicators of warm climate) and a decline of Alpine steppes (indicators of cold climate) at different time points including the GLGM [75, 76] suggesting that vegetation communities did not support the distribution of pikas at this time points given the mismatch in habitats.

### Population responses of pikas on the Indo-Tibetan plateau

Our PSMCs largely track the glacation history of the CBP with the largest population sizes of pikas either being at LIG *(O. nubrica)* or MIS-6 (*O. ladacensis* and *O. macrotis*) before crashing into the Holocene, much like other range restricted pikas such as *O. sikkimaria* [50] . Such disparity in the timing of population peaks between species might have arisen either due to *O. nubrica* showing delayed effective population size responses to changing climate [77, 78] or the less likely but incorrect choice of mutation rates and generation times while scaling plots (Figure S2). Species on the Tibetan plateau have typically experienced population size increases dating back to the MIS-6 period, and includes domestic and wild mammals [47, 49], birds [68, 74] and reptiles [79]. Of the limited studies that have examined demographic history through PSMCs on the Tibetan plateau, patterns recorded for the Iberian Ibex [45] and the Tibetan Wolf [46] show contrasting patterns. Given the absence of any glaciation events post MIS-3 and MIS-4 (which was also very mild on the CBP), pikas are likely to have remained isolated on cool mountain tops causing them to inbreed for long periods, leading to a decline in population size in the absence of out-breeding rescue [50, 58, 59].

Recent studies of the subgenus *Ochotona* have suggested that outbreeding and adaptive introgressions between *O. curzoniae* and other members of the sub-clade have rescued populations over historic timescales [50]. While recent inbreeding is known to affect the estimation of coalescence times and Ne [80, 81], inbreeding at older time scales doesn’t necessarily affect these estimates [82, 83]. We, therefore, believe that the population demographic signatures obtained by us across species are driven by the lack of severe glaciation events in the CBP in the last 1mya and inbreeding depression post the LIG. This requires further validation through other analyses of formal runs of homozygosity analysis (ROH) [80, 82].

## Conclusion

While our PSMCs are likely to be true depictions of population responses of pikas from the Indo-Tibetan plateau (Changthang Biotic Province), our SDMs are capturing the general patterns of species responses (elevational shift) across their geographic range. We find that lifestyle (burrowing and rock-dwelling), did not interact with population trajectories recovered. We hypothesize that pikas in the Changthang Biotic Province had the largest population sizes during the cool period of MIS-6, were forced up the elevational gradient during the warm period of LIG and remained isolated and inbred in the absence of a glaciation rescue causing the drop in effective population sizes. Sampling demographics of cold-adapted specialists on the Indo-Tibetan plateau in a spatially explicit manner with consideration of local geology should shed more insights for conservation management of these species under threat.

## Methods

### Species distribution modeling (SDMs)

Occurrence records: We collected primary points of occurrence of all three species of pikas between 2017 and 2023 in the Tibetan Marginal Mountains, Trans-Himalayas, Ladakh, India. We supplemented this data with secondary points from published literature, GBIF (Global Biodiversity Information Facility), iNaturalist and museum records. Only confirmed observations on citizen science platforms with a photo where species identification could be verified were used. Our dataset included 63 occurrence points for *O. ladacensis,* 203 points for *O. macrotis,* and 93 points for *O. nubrica*.

### Raster data processing

Raster layers of bioclimatic variables (Bio1 - Bio19) were sourced from Bioclim [84] at 1km resolution for four different time slices: LIG (120 kya), LGM (20 kya), Mid-Holocene (5000 ya) and current [85]. While the LIG, Mid-Holocene, and current data were available at a 30-second resolution, the LGM data was available only at a coarser, 2.5-minute resolution. The LGM data was interpolated and downscaled to a 30-second resolution to match other raster datasets. A conservatively large area (coordinates - 62.2749, 25.5666: 104.9499, 46.8416) encompassing the distribution of the most widely distributed species was chosen as the study area. (Figure 3). The advanced digitizing toolbox was used to create this shape file, and all rasters were clipped to this mask. While doing so, all rasters were converted to the same projection system (EPSG 4326: WGS 84), with no data pixels being set to -9999 and converted to an ‘ASCII’ file format as required by MAXENT. Processed rasters were imported into QGIS, and the spatial extent, pixel sizes, and range of pixel values were inspected before further downstream processing. The ENMevaluate function was run for occurrence data at current timescales to determine the best model settings for MAXENT. Model performance under different parameter settings was evaluated for each species using AIC scores, and the best model settings for each species are presented (Table S1). Occurrence records were spatially thinned (1 point/km) to avoid data over-fitting. Post spatial thinning, the dataset had 33 points for *O. ladacensis*, 37 points for *O. nubrica*, and 121 points for *O. macrotis*. A bias file was also created to account for sampling biases by modifying existing R scripts (Josh Banta Lab, https://sites.google.com/site/thebantalab/tutorials#h.zsfviftgr2).

### Genome sequencing and De-novo assembly

One tissue sample was collected for each of the three species using standard trapping methods with baited Tomahawks or from opportunistic roadkills. These tissues were stored in ethanol at -20°C for downstream processing. DNA was extracted from tissues using QIAGEN Blood and Tissue Kits with a modified protocol to include an additional digestion step with AL buffer for 20 minutes at 75°C and a digestion with RNase. Extracted DNA was quantified using a Qubit fluorometer and sequenced using an Illumina platform sequencer at 30x coverage. The reads obtained for each sample were subject to quality checks using FASTQC [86] and trimmed with a quality score cutoff of 30 using TRIM GALORE [87]. Trimmed reads were repaired and quality checked before de-novo assembly using the Megahit [88] pipeline under default settings.

### Genome quality assessment

Quality and completeness of genomes assembled using Megahit were assessed using BUSCO [89] against the mammalian database using Augustus [90]. Proteins predicted were merged together for each species, prefixed with species identifiers and prepped for comparative genomic analyses. Orthovenn3 [91, 92] was used to identify orthologous gene clusters across the draft genomes assembled. It does so by employing BLASTP in conjunction with constructing a similarity graph followed by clustering this network [91, 92].

### Pairwise sequential Markovian Coalescent (PSMC) analysis

While demographic inferences are robust to genome assembly quality [78], the presence of repeats in mammalian genomes, mitochondrial DNA, and sex chromosomes are known to affect demographic inference [32, 82]. Assemblies were therefore, hard masked for repeats using RepeatMasker [93], following which, contigs were mapped to reference mitochondrial genomes of *O. princeps* (NCBI Accession number: NC_005358.1) and sex chromosomes of *O. cuniculus* (NCBI Accession number: NC_067395.1) using BLAST to identify and remove such contigs in assemblies. Trimmed reads from the Illumina sequencing experiment were marked for PCR duplicates and mapped back to this assembly to generate a binary alignment file (.bam) using ‘samtools mpileup’ [94]. A ‘diploid.fq’ file was generated following standard PSMC protocols for downstream processing using default depth cutoffs when variant calling. The fq2psmca tool was used to generate psmcfa files, and PSMCs were performed with 100 bootstraps under settings that detected more than 1000 recombinations at each of the timesteps (ON: -N25 -t5 -r5 -p "26*2+4+7+1"; OL: -N25 -t15 -r5 -p "4+25*2+4+6"; OM: -N25 -t9 -r5 -b -p "26*2+4+7+1") after post ensuring that the average depth of sequencing was >30x [27, 30] (Supplementary Figures 1).

The psmc_plot.pl function was used to scale PSMCs and generated text output for plotting bootstraps on R. While scaling plots, the mutation rate for all species was set at 2.96*10^-9^ following published literature on pikas and rodents (mean mutation rate for nuclear protein-coding genes) [50, 95, 96]. Different generation times were used for different species of pikas based on both field guides and natural history data collected while on the field between 2019-2023 (*O. nubrica* - 0.5 years, *O. ladacensis* - 1 year, *O. macrotis* - 1 year) after scaling them by 2 [11, 32]. We additionally modeled the sensitivity of the PSMC results obtained using various parameters of mutation rates and generation times (Figure S2).

## Acknowledgments

We thank the Jammu and Kashmir Wildlife Department and Department of Wildlife Protection, Ladakh for permits to collect small tissue samples from caught animals/ roadkills (471-75/WLWL/E.P/2022-23) for whole genome sequencing, Senan D’Souza & Dhanesh Ponnu for help with trapping animals and mapping distributions of species in Ladakh, Dr. Meghana Natesh, Dr. Nivetha Murugesan and Vinay KL for comments on the work, and IISER Tirupati for funding and logistics.

## Author contributions

HK - conceptualization, analysis, writing, sample collection; NV - conceptualization, inputs on analysis, reviewing the manuscript; NR - conceptualization, reviewing the analysis, reviewing the manuscript, sample collection.

