## Supplementary File for "Asian pika populations track local glaciation events through the Pleistocene"

| Variable | Percent contribution | Permutation importance |
|---|---|---|
| bio9_mean_temp_dri_qtr | 28.8 | 27.4 |
| bio15_precip_seasonality | 17 | 36.1 |
| bio7_temp_ann_rang | 11 | 0 |
| bio17_precip_dri_qtr | 8.6 | 7.2 |
| bio8_mean_temp_wet_qtr | 7.8 | 22.7 |
| bio19_precip_cold_qtr | 7.3 | 5.9 |

Table S1: Top variables (Percent contribution > 5%) identified in MAXENT models of *Ochotona nubrica* (ON).

| Variable | Percent contribution | Permutation importance |
|---|---|---|
| bio15_precip_seasonality | 29.3 | 8.4 |
| bio10_mean_temp_wrm_qtr | 26.8 | 0 |
| bio8_mean_temp_wet_qtr | 13 | 30.8 |
| bio14_precip_dri_mnth | 10.2 | 0.8 |
| bio3_isothermality | 7.8 | 13.4 |
| bio9_mean_temp_dri_qtr | 7 | 22 |

Table S2: Top variables (Percent contribution > 5%) identified in MAXENT models of *Ochotona ladacensis* (OL).

| Species | Timescale | Average elevation (m) |
|---|---|---|
| *Ochotona ladacensis* | Current | 4388.49 |
| | LGM | 1900.53 |
| | LIG | 1506.56 |
| *Ochotona nubrica* | Current | 4444.35 |
| | LGM | 1700.75 |
| | LIG | 552.95 |
| *Ochotona macrotis* | Current | 3091.86 |
| | LGM | 2321.85 |
| | LIG | 3057 |

Table S3: Elevational distribution of pikas at historical and current timescales based on niche suitability (> 0.4).

| Variable | Percent contribution | Permutation importance |
|---|---|---|
| bio1_ann_mean_temp | 26.7 | 27.7 |
| bio14_precip_dri_mnth | 18.9 | 0.6 |
| bio9_mean_temp_dri_qtr | 16.5 | 11.8 |
| bio15_precip_seasonality | 15.9 | 21.4 |
| bio11_mean_temp_cold_qtr | 6.6 | 0 |

Table S4: Top variables (Percent contribution > 5%) identified in MAXENT models of *Ochotona macrotis* (OM).

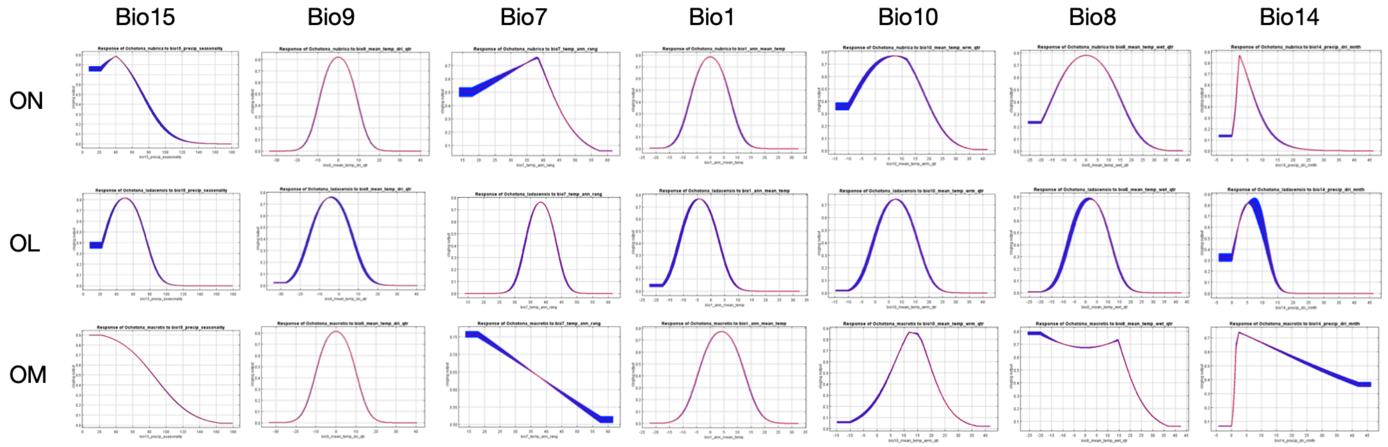


Figure S1: Uni-variate response curves of variables important in driving species occurrence at time scales. Variables important for each species were pooled together and are represented here.

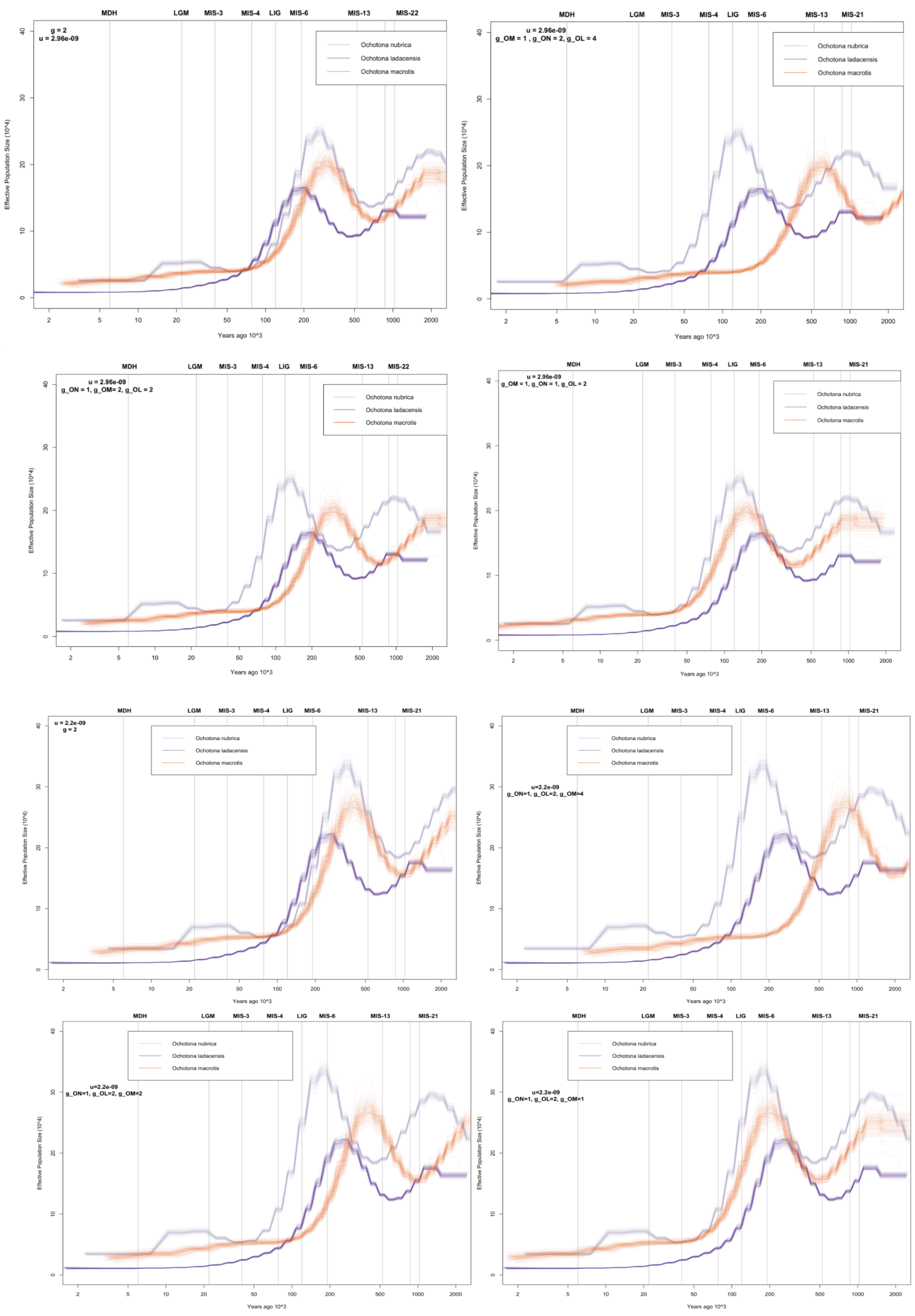


Figure S2: Sensitivity of PSMC to scaling by changing both mutation rates and generation times.
